# Balancing selection and the crossing of fitness valleys in structured populations: diversification in the gametophytic self-incompatibility system

**DOI:** 10.1101/2021.11.20.469375

**Authors:** Roman Stetsenko, Thomas Brom, Vincent Castric, Sylvain Billiard

## Abstract

The self-incompatibility locus (*S*-locus) of flowering plants displays a striking allelic diversity. How such a diversity has emerged remains unclear. In this paper, we performed numerical simulations in a finite island population genetics model to investigate how population subdivision affects the diversification process at a *S*-locus, given that the two-genes architecture typical of *S*-loci involves the crossing of a fitness valley. We show that population structure slightly reduces the parameter range allowing for the diversification of self-incompatibility haplotypes (*S*-haplotypes), but at the same time also increases the number of these haplotypes maintained in the whole metapopulation. This increase is partly due to a higher rate of diversification and replacement of *S*-haplotypes within and among demes. We also show that the two-genes architecture leads to a higher diversity in structured populations compared with a simpler genetic architecture where new *S*-haplotypes appear in a single mutation step. Overall, our results suggest that population subdivision can act in two opposite directions: it renders *S*-haplotypes diversification easier, although it also increases the risk that the self-incompatibility system is lost.

## Introduction

Adaptation often occurs on rugged fitness landscapes with multiple peaks and valleys (*e*.*g*. Korona et al. 1994; Burch and Chao 2000; Weinreich et al. 2006; Kvitek and Sherlock 2011; Salverda et al. 2011; Vos et al. 2015, for a review see de Visser and Krug 2014). This is especially true when the trait is genetically complex and the emergence of evolutionary novelties is conditioned on the beneficial combination of epistatic mutations in a single genome (Whitlock et al., 1995). In many cases, attaining a higher fitness peak can require a succession of mutations where some are deleterious on their own. Under the “shifting balance theory” popularized by Wright (1932) this process is referred to as the crossing of fitness valleys. The rate at which populations move across the fitness landscape and eventually cross fitness valleys depends on a subtle balance between mutation rate, population size and the distribution of mutational effects (*e*.*g*. Weissman et al. 2009; Bovier et al. 2019 in asexual panmictic populations), in combination with genetic exchange across space (*e*.*g*. Bitbol and Schwab 2014; Komarova et al. 2014; McLaren 2016 in asexual spatially divided populations) or between genomes (the recombination rate in sexual populations, *e*.*g*. Weissman et al. 2010). Overall, the rate at which fitness valleys are crossed depends on the time taken for the epistatic mutations to occur in the same genome, and the probability of fixation and time of residence of mutations in the population. Theoretical studies investigating the conditions under which fitness valleys can be crossed have mostly assumed directional selection, so their predictions do not apply to all traits. Many complex genetic systems, in particular, evolve under balancing selection, including in particular pathogens escaping the immune system of their host (Bowen and Walker, 2005; Fernandez et al., 2005; Shih et al., 2007), or the genes involved in avoidance of self-fertilization in fungi, plants or ciliates (Billiard et al., 2011). Studying how fitness valleys are crossed in such systems is particularly relevant because they are often characterized by a very large allelic diversity at superloci, suggesting a high rate of emergence of evolutionary novelties and thus frequent fitness valley crossing.

The self-incompatibility (SI) system in Angiosperms is particularly relevant to address this question. SI is present in about 40% of Angiosperm families, where it allows recognition between pollen and pistils for selfing avoidance (Igic et al., 2008; McCubbin and Kao, 2000; Barrett, 2002). SI is often determined by a single locus, called the *S*-locus. The *S*-locus is characterized by high haplotypic diversity which is maintained by a particular type of balancing selection called negative frequency-dependent selection, whereby individuals expressing rare haplotypes can mate with more individuals than the ones carrying frequent haplotypes (Wright, 1939). A series of early theoretical works approximated the number of alleles that can be maintained at the *S*-locus (*S*-alleles) in a population under various conditions (*e*.*g*. population size, mutation rate, deleterious mutations) assuming that *S*-alleles are generated by mutations occurring at the *S*-locus at a given rate (Wright, 1939, 1964; Yokoyama and Hetherington, 1982; Muirhead, 2001; Uyenoyama, 2003). However, these models ignored the mechanisms by which new *S*-alleles can emerge, and more recent works focused on the conditions of emergence of new *S*-haplotypes under a variety of more realistic assumptions (Uyenoyama et al., 2001; Gervais et al., 2011; Sakai, 2016; Bod’ová et al., 2018; Harkness et al., 2019), in particular through an evolutionary step implying the crossing of a fitness valley.

In this study we focused on gametophytic SI (GSI), in which the pollen specificity is determined by its own haploid genome and *S*-alleles are codominant in pistils. The genetic architecture of the *S*-locus is typically relatively simple, with male and female specificities encoded by distinct but tightly linked genes (often one each, Fujii et al., 2016, but see non-self recognition systems, *e*.*g*. Bod’ová et al., 2018). Uyenoyama et al. (2001) studied the conditions of emergence of a new *S*-haplotype by mutations on both genes. They considered that new *S*-haplotypes are formed first by a mutation on the male or the female determinant, creating an intermediate haplotype with a new pollen or pistil specificity, but no matching recognition at the other gene. This haplotype is thus self-compatible (SC) and its selfed offspring will suffer from inbreeding depression. This intermediate haplotype thus represents a fitness valley that needs to be crossed in order to create a new *S*-haplotype. This happens when the first mutation is followed by a compensatory mutation on the female or on the male determinant (depending on the nature of the first mutation), restoring a fully functional new *S*-haplotype in which the pistil specificity matches that of the pollen. Uyenoyama et al. (2001) showed that in an infinite population, new *S*-haplotypes can emerge for high values of inbreeding depression and low to intermediate values of the rate of self-pollination. In these conditions, the intermediate SC haplotype invades when it is rare, but it does not go to fixation, allowing the second mutation to arise and create the new *S*-haplotype. However, they also showed that in the course of this process, the ancestral *S*-haplotype was most often lost by competitive exclusion by its mutated SC haplotype. Thus, this model predicted that even though new *S*-haplotypes can emerge, the new derived *S*-haplotypes tend to replace their ancestral copies along the process, such that in the end the overall number of *S*-haplotypes is expected to remain constant. The conditions under which diversification of *S*-haplotypes is possible are thus restricted either because the SC intermediate haplotype cannot invade at all or because its ancestor is lost. In this model, crossing fitness valleys thus results in *S*-haplotypes turnover rather than diversification (Box 1, Fig. S1).

### Box 1

The question of *S*-haplotype emergence is the following: how is it possible to go from *n* to *n* + 1 *S*-haplotypes segregating in a population, given an intermediate SC haplotype which can have a fitness lower or higher than its ancestral (*S*_*n*_) and derived new (*S*_*n*+1_) *S*-haplotypes depending on the cost of inbreeding depression relative to the strength of negative frequency dependent selection (Fig. S1A, B and C).

The number of *S*-haplotypes increases when a fitness valley is crossed either because the SC intermediate haplotype has a fitness lower than both *S*_*n*_ and *S*_*n*+1_ (Fig. S1B), or because *S*_*n*+1_ has a higher fitness than *S*_*n*_ when it co-occurs with the SC intermediate haplotypes and thus *S*_*n*_ is lost (Fig. S1C) Uyenoyama et al. (2001) and Gervais et al. (2011)) showed that fitness valley crossing can be achieved by two mechanisms: either because the ancestral *S*-haplotype *S*_*n*_ and the intermediate SC haplotype can stably coexist under negative frequency dependent selection, or because haplotypes with the lowest fitness can be maintained in the population long enough because of genetic drift. Genetic drift is thus an important mechanism allowing fitness valley crossing under either directional (Weissman et al., 2009) or negative frequency dependent (Gervais et al., 2011) selection.

Gervais et al. (2011) further analyzed Uyenoyama et al. (2001)’s model, but in a finite population with recurrent mutations. They showed that the rate at which new *S*-haplotypes emerge depends on a balance between population size, mutation rates and selection strength (inbreeding depression cost *vs*. negative frequency-dependent selection). More importantly, *S*-haplotype diversification occurred under a relatively wide range of parameters because the residence time of SC haplotypes can be long in finite populations. They further showed that diversification is a self-attenuating process, whereby the increase in the number of *S*-haplotypes in the population renders the emergence of novel *S*-haplotypes in turn less likely. As a result, regardless of the parameter values considered, the maximal number of *S*-haplotypes maintained at stationary state is predicted to rarely go beyond a dozen, almost an order of magnitude lower than the observed haplotypic diversity in natural populations (from a few dozens to *>* 100 observed *S*-haplotypes (Lawrence, 2000; Castric and Vekemans, 2004)). Hence, current models in finite or in infinite isolated populations fall short in explaining the entire diversity encountered at the *S*-locus in natural populations (but see Sakai, 2016).

Wright (1939) was the first to propose that population subdivision might contribute to the large diversity of *S*-haplotypes, but he did not account for the particular two-genes genetic architecture underlying SI. Uyenoyama et al. (2001) and Gervais et al. (2011) latter studied diversification of *S*-haplotypes with the two-genes genetic architecture but they did not explore the effect of population subdivision. Predicting the effect of population sub-division on the diversification process is not straightforward as population subdivision can have opposite effects on the maintenance of *S*-haplotypes at the global (metapopulation) or local (deme) scale because: 1) *S*-alleles can be lost by chance in isolated demes (genetic drift), 2) balancing selection increases the invasion probability of new *S*-alleles when the local *S*-allele diversity is low, and 3) dispersal homogenizes *S*-allele diversity among demes (Schierup, 1998; Schierup et al., 2000; Muirhead, 2001). In addition, population subdivision facilitates the breakdown of SI because it decreases the local effective number of *S*-haplotypes helping the invasion of SC haplotypes in the metapopulation (Brom et al., 2020). How population subdivision would globally affect diversification thus remains unclear.

In this paper, we used population genetics simulations to evaluate the effect of population structure on the dynamics of *S*-haplotypes diversification. First, we compared the parameter range allowing *S*-haplotypes to diversify in structured populations with that in unstructured populations. Second, we determined how the number of *S*-haplotypes maintained at steady state varies with the degree of population structure. Third, we disentangled the mechanisms underlying the effect of population structure on *S*-haplotypes diversification. Finally, in order to study the more general issue of fitness valley crossing in structured populations, we investigated the interaction between population structure and the underlying genetic architecture of the *S*-locus.

## Models and Methods

### Genetic and demographic model

A subdivided population of diploid hermaphroditic plants is assumed with a gametophytic SI mating system. The SI phenotype, *i*.*e*. the expressed specificities in pollen and pistils, is determined by the *S*-haplotypes of the individuals. Mating can only occur between individuals expressing different pollen and pistil specificities. Pollen specificity is determined by the haploid genome of the gametophyte (the pollen grain), and the pistil specificity is codominantly determined by the diploid genome of the sporophyte (the adult plant). *S*-haplotypes are composed of two completely linked genes: the pistil-part gene, denoted *A*, determines the female specificity; the pollen-part gene, denoted *B*, determines the male specificity. We denote *S*_*i*_ an haplotype expressing specificity *i* at the pollen and pistil genes, and *A*_*i*_ and *B*_*i*_ the alleles present at genes *A* and *B*, respectively. Hence, an individual with genotype *S*_*i*_*S*_*j*_ (carrying haplotypes *A*_*i*_*B*_*i*_*/A*_*j*_*B*_*j*_) is self-incompatible since its individual pollen grains express either the specificity *i* or *j*, and its pistils express both specificities *i* and *j*.

An analytical model taking into account diversification of *S*-haplotypes with the two-genes genetic architecture and population subdivision was not possible to build. Indeed, the rate of diversification of *S*-haplotypes depends on the probability that a compensatory mutation arises in a population of SC haplotypes. Evaluating this probability would have implied the construction of a stochastic model to compute the average residence time of those SC haplotypes in the population, taking into account the effect of drift, inbreeding depression and compatibility between haplotypes. Because, to our knowledge, such a model is not mathematically tractable we resorted to individual-based simulations of a finite island model with non-overlapping generations. Initially, the population was composed of self-incompatible individuals with heterozygous *S*-genotypes randomly drawn among the *n* possible *S* haplotypes. The metapopulation was divided into *p* equally-sized demes of *N* individuals. Pollen dispersed at rate *d*_*p*_ between demes (we neglected seed dispersal). During the life cycle, events occurred in the following order: 1) Mutations at genes *A* and *B*; 2) Pollen migration between demes; 3) Pollination and fertilization; 4) Demographic events: reproduction, adults death and recruitment.

### Mutation

We assumed a maximum of *k* possible specificities in the population (1 ≤ i ≤ *k*). In other words, we considered a *k*-alleles mutation model at both the pollen and the pistil genes. Mutations are recurrently introduced into the population at the *S*-locus. Under the 2-steps mutation model (Fig. 1B), mutations occur with probability *µ*_2_, on the pollen gene (*B*_*i*_ → *B*_*j*_) or on the pistil gene (*A*_*i*_ → *A*_*j*_), independently. A mutation randomly changes a specificity *i* into a specificity *j ≠ i* (1 ≤ *j* ≤ *k*), drawn from a uniform distribution. In line with the notation of Uyenoyama et al. (2001), a mutation can thus give rise to four different haplotypes, depending on the ancestral haplotype and the modified gene: 1) a pollen-part mutant self-compatible (SC) haplotype, denoted *S*_*b*_; 2) a pistil-part mutant SC haplotype, denoted *S*_*a*_; 3) an already existing *S*-haplotype, denoted *S*_*n*_; 4) a new *S*-haplotype, denoted *S*_*n*+1_ (1 ≤ *n* + 1 ≤ *k*).

**Fig. 1:**
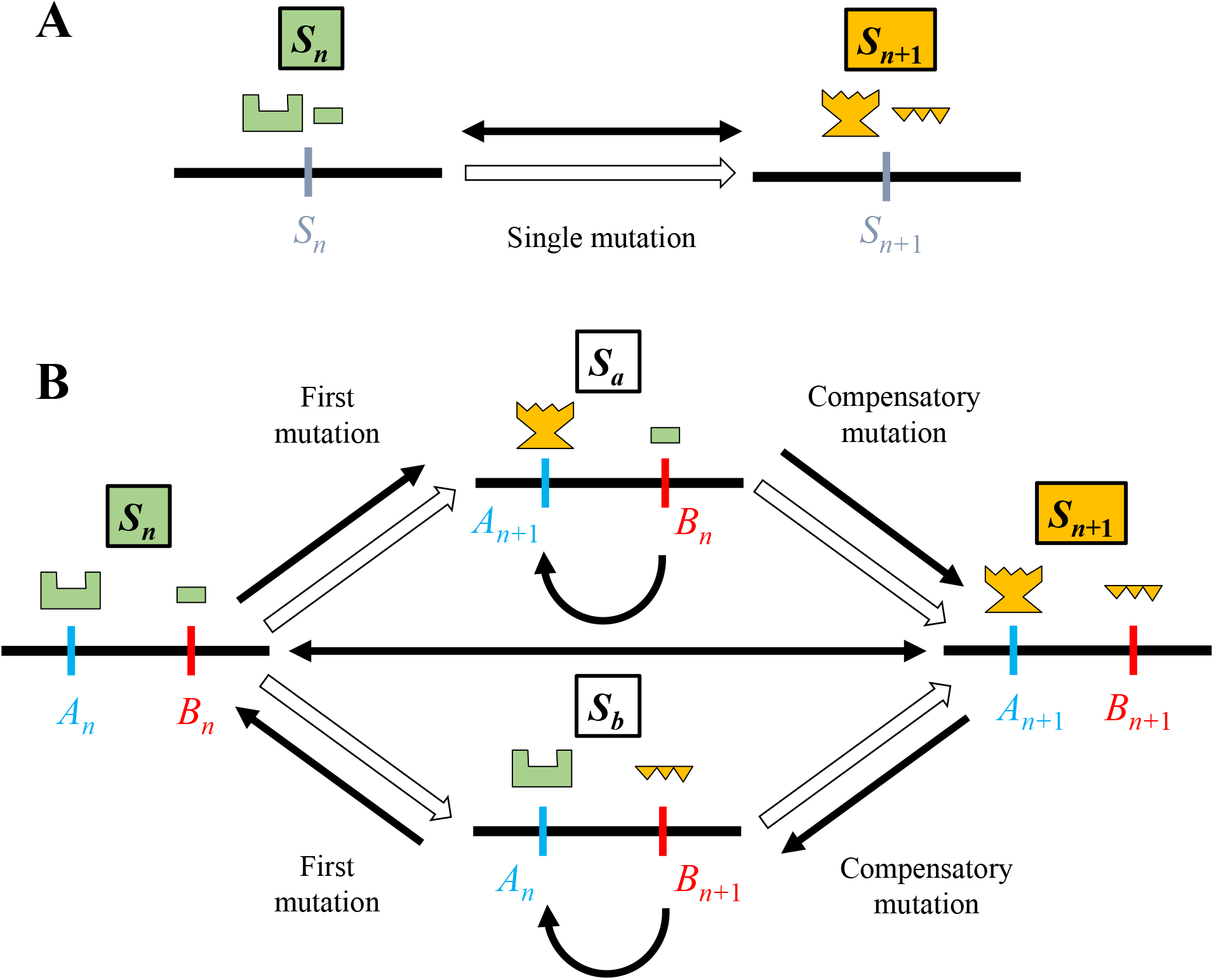
Mutational pathway by which new haplotypes are created in the 1-step mutation model (A) and in the 2-steps mutation model (B; Uyenoyama et al., 2001; Gervais et al., 2011). Horizontal black lines represent chromosomes carrying the *S*-haplotype which is composed of just one gene in the 1-step mutation model and of two genes in the 2-steps mutation model with gene *A* as the female determinant (blue vertical lines) and gene *B* as the male determinant (red vertical lines). *S*_*n*_ is the ancestral *S* haplotype, *S*_*n*+1_ the new *S* haplotype, *S*_*a*_ the pistil-part mutant SC haplotype and *S*_*b*_ the pollen-part mutant SC haplotype. Incompatible specificities are represented by matching shapes and identical colours. White arrows represent mutation steps and black arrows indicate that the pollen carrying the haplotype at the base of the arrow can pollinate the pistil carrying the haplotype at the tip of the arrow. Double black arrows indicate mutual compatibility between haplotypes. Modified from Uyenoyama et al. (2001).

The pistil- and pollen-part mutant haplotypes *S*_*a*_ and *S*_*b*_ are self-compatible because their pollen and pistils express different specificities. Note that haplotypes *S*_*a*_ and *S*_*b*_ can express a specificity *n* +1 which is not already present in the population. A new *S*-haplotype *S*_*n*+1_ is formed when an ancestral haplotype has undergone two mutations: a first mutation giving a SC haplotype (*S*_*b*_ or *S*_*a*_) and a second compensatory mutation restoring the *S*-haplotype function.

In order to compare the impact of the genetic architecture of *S*-haplotypes on the diversification dynamics, we also considered a 1-step mutation model. In this model, haplotype *S*_*i*_ mutates in a single step to *S*_*j*_ at rate *µ*_1_ per haploid genome per generation (*i*.*e*. both alleles *A*_*i*_ and *B*_*i*_ simultaneously mutate to alleles *A*_*j*_ and *B*_*j*_). *j* is randomly and uniformly chosen such that *j* ≠ *i* and 1 ≤ *j* ≤ *k*. This 1-step model, where the *S*-locus is not split into a female and a male genes and mutations do not generate self-compatible haplotypes, is the one classically considered in models on the evolution and maintenance of GSI (*e*.*g*. Wright, 1939; Schierup, 1998; Gervais et al., 2014; Brom et al., 2020).

### Pollination

Each individual of the next generation is formed by randomly drawing a female and a male gamete. Firstly, a female is drawn in the local deme (no seed dispersion) according to its relative fitness in the population, which differs between selfed and outcrossed individuals because of inbreeding depression *δ*. The fitness of selfed *vs*. outcrossed individuals is *W*_*self*_ = 1 − *δ* and *W*_*out*_ = 1 respectively (the probability that an individual is chosen as a mother is thus weighted by its own fitness relative to all the other individuals). Once a mother is chosen, an ovule is produced by randomly choosing one of the two chromosomes of the mother, with equal probability. Secondly, a father is drawn from the same deme as the mother with probability 1 – *d*_*p*_ or from a different deme with probability *d*_*p*_. The probability for a plant to be chosen as a father is also weighted by its own fitness because of inbreeding depression (*W*_*self*_ = 1 − *δ* if inbred, *W*_*out*_ = 1 if outcrossed). If the pollen grain is not compatible with the pistil of the chosen mother, another pollen is randomly drawn until fertilization can occur (hence all chosen ovules eventually find a compatible pollen, precluding pollen limitation). In SC individuals, compatible pollen can come either from their own pollen or from other individuals and the probability that an SC individual reproduces by selfing corresponds to the ratio of its own compatible pollen over the compatible pollen present in the total pollen pool. The effective selfing rate of an individual *i* is thus given by

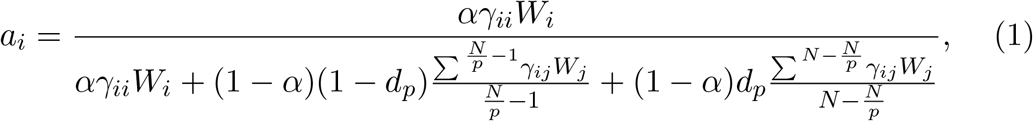

where *α* is the proportion of self-pollen, *γ*_*ii*_ the number of SC haplotypes carried by individual *i, γ*_*ij*_ the number of compatible haplotype an individual *j* has with an individual *i* (*γ*_*ii*_ and *γ*_*ij*_ can thus be 0, 1 or 2), *W*_*i*_ the pollen production of individual *i* and *W*_*j*_ the pollen production of individual *j* (*W*_*i*_, *W*_*j*_ = *W*_*self*_ or *W*_*out*_ depending on whether individuals *i* and *j* are selfed or outcrossed). The numerator and first term of the denominator correspond to compatible self pollen of individual *i*, the second term of the denominator corresponds to compatible pollen of other individuals from the same deme and the third term of the denominator corresponds to compatible pollen of individuals from other demes. The outcrossing rate of an individual *i* is thus 1 − *a*_*i*_.

### Verification of the simulations

We first verified the validity of our simulation program by comparing its outcomes with previous results from Gervais et al. (2011) and Uyenoyama et al. (2001) for the 2-steps mutation model in a panmictic population, and from Schierup (1998) for the 1-step mutation model in a subdivided population with the same parameter values. We recovered results similar to Gervais et al. (2011) and Uyenoyama et al. (2001) (Figure S2 and Figure S3) when comparing (i) the range of parameters values where allelic diversification occurrs in a panmictic population of size *N* = 5000; (ii) the probability of diversification in 100 replicated runs; and (iii) the dynamics of SI and SC haplotypes. We also recovered results similar to Schierup (1998) with our 1-step mutation model when comparing the average number of haplotypes at the scale of the whole metapopulation or of a single deme (*<* 3% differences, Figure S4), for identical parameter values and the same estimation method.

### Diversification probability

We first studied how dispersal affects the probability of diversification. Simulations were repeated a hundred times for each parameter set (inbreeding depression *δ*, self-pollination rate *α*, pollen dispersal rate *d*_*p*_, mutation rate *µ*_2_, number of demes *p* and size of the demes *N*). Simulations were started with *n* = 5 functional *S*-alleles and no self-compatible haplotypes. The number of possible haplotypes was set to *k* = 20. Simulations were run until one of the three following events occurred (note that since haplotypes could appear by mutation and be lost by drift when rare, we arbitrarily took into account only haplotypes with a frequency higher than 0.01): i) *loss of SI* : when SC mutant haplotypes had invaded the population and all *S*-haplotypes were lost; ii) *diversification*: when the number of *S*-haplotypes reached *n* + 1 = 6 and the frequency of SC haplotypes was lower than 0.1; iii) *undecided* : When none of the two previously described events happened before 5.10^5^ generations. We estimated the probability of diversification by the number of runs where the event *diversification* occurred among the one hundred runs performed for a given parameter set.

### Number of *S*-haplotypes at stationary state

We then studied the effect of population structure on the number of *S*-haplotypes maintained at equilibrium in the metapopulation. Because we wanted to disentangle the effect of population structure on the number of *S*-haplotypes maintained at steady state from its effect on the probability of diversification, we ran simulations using values of *d*_*p*_ and *µ*_2_ under which the results above indicated that *diversification* occurred with probability ≥ 0.5 (Fig. S5). Otherwise the number of *S*-haplotypes at steady state would mostly reflect the probability of diversification. For the same reason we also discarded simulation runs where SI was lost or no diversification had occurred. Simulations were run as described in the previous section one hundred times for a given parameter set, with the exception that we let the metapopulation evolve for an arbitrarily large number of generations (10^6^) instead of stopping it when a diversification occurred. The average number of *S*-haplotypes at steady state 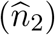 was estimated over all replicates for a given parameter set by calculating the number of *S*-haplotypes with a frequency *>* 0.01 during the last 50,000 generations.

### Interaction between the genetic architecture and population structure

In order to evaluate the effect of the genetic architecture of SI on diversification in a structured population, we compared the average number of *S*-haplotypes at steady state under the 1- *vs*. 2-steps mutational models. For this comparison to be meaningful, the number of *S*-haplotypes generated at equilibrium in a non-structured population with the same mutation rate must be the same in both models. In the 1-step mutation model, *µ*_1_ is the rate at which new *S*-haplotypes. In the 2-steps mutation model, this rate is not straightforward since it depends on the mutation rate of the pistil and pollen genes *µ*_2_ but also on the frequency of SC mutants that are the substrate for the compensatory mutations (Gervais et al., 2011). We defined an *effective mutation rate* as the value *µ*_1_ such that the average number of *S*-haplotypes is similar in the 1- *vs*. 2-steps mutation model (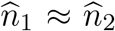 *i*.*e*. the value of *µ*_1_ that gave the closest value of 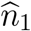 from 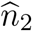) in a non-subdivided population. We then ran simulations under both mutational models for the same parameter sets, including the effective mutation rate, for different levels of population structure. Comparing the number of *S*-haplotypes at steady state under both mutational models allowed us to isolate the specific effect of the interaction between population structure and genetic architecture.

### Scenarios of fitness valleys crossing in structured populations

Population structure expands the number of scenarios by which new *S*-alleles can be formed, according to whether those haplotypes arose in the same or different demes and whether the intermediate SC haplotype was maintained along with other SI haplotypes in the different demes. We identified five different possible scenarios (summarized in Figure 2):

1. Local diversification — The two mutations occurred in the same deme and the ancestral haplotype was not lost from this deme. Under this scenario, the number of *S*-haplotypes increases at the deme scale, and consequently also at the metapopulation scale.
2. Local replacement— The two mutations occurred in the same deme but the ancestral haplotype was lost from this deme. Under this scenario, the number of *S*-haplotypes does not change at the deme scale, but it increases at the metapopulation scale (provided that the ancestral haplotype is present in at least one other deme).
3. Allodiversification — The two mutations occurred in different demes (*i*.*e*. at least one dispersal event of the intermediate SC haplotype occurred before the compensatory mutation) and the ancestral haplotype was conserved in the deme where the compensatory mutation occurred. Under this scenario, the number of *S*-haplotypes increases at the deme scale, and consequently also at the metapopulation scale.
4. Alloreplacement — The two mutations occurred in different demes (*i*.*e*. at least one dispersal event of the intermediate SC haplotype occurred before the compensatory mutation) but the ancestral haplotype was lost from the deme where the compensatory mutation occurred. Under this scenario, the number of *S*-haplotypes does not change at the deme scale, but it increases at the metapopulation scale.
5. Global replacement — The two mutations occurred in different demes but the ancestral haplotype was lost from the whole metapopulation. Under this scenario, the number of *S*-haplotypes does not change neither at the deme scale nor at the metapopulation scale.

**Fig. 2:**
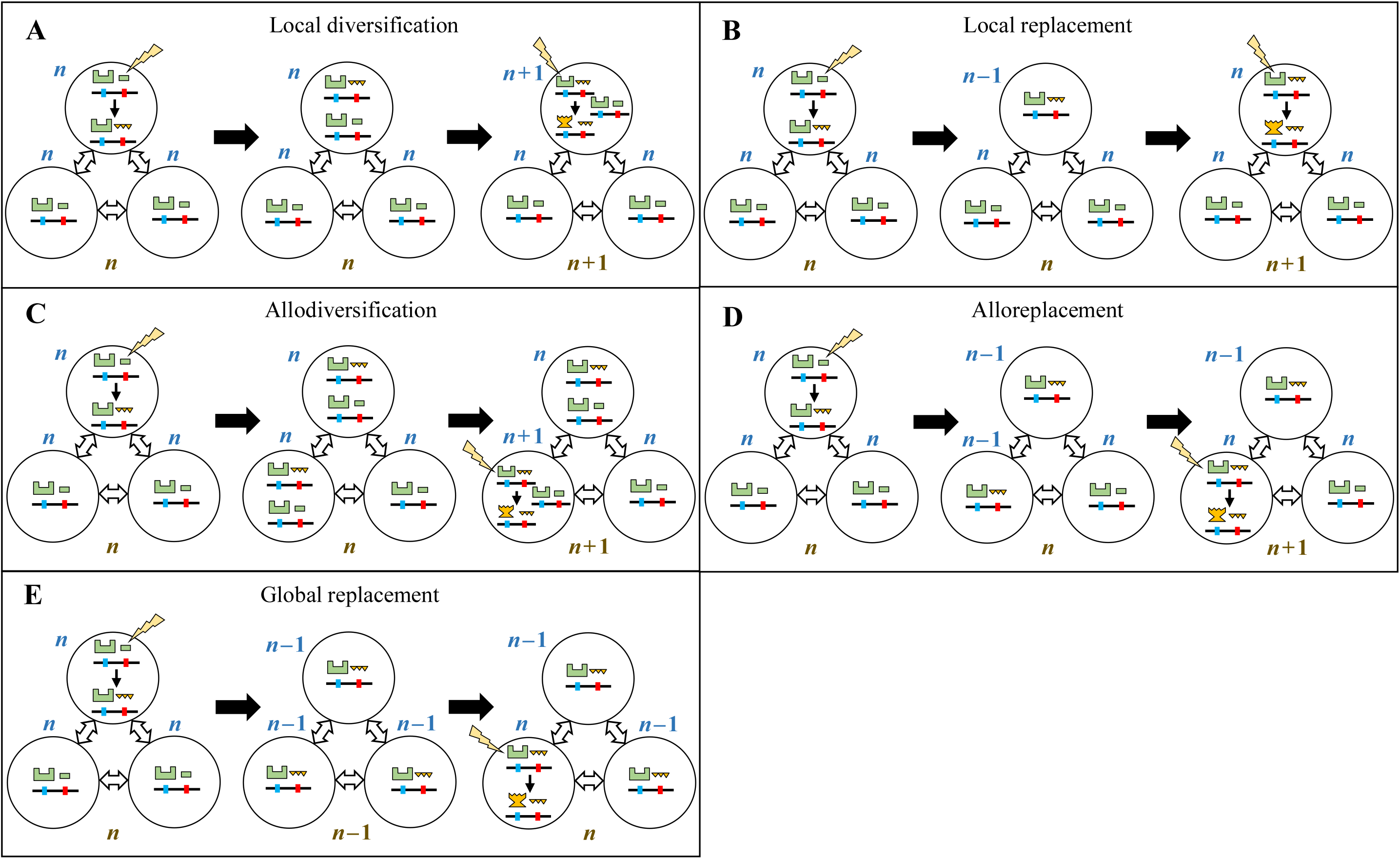
Five different scenarios for the emergence of a new *S*_*n*+1_ *S*-haplotype in a subdivided population. Note that only one allelic line undergoing diversification is depicted although it is segregating along with many others in the population. Three demes are depicted, with different possible *S*-haplotypes (pistil gene on the left, pollen gene on the right, the lightning bolt depicts mutation; see Figure 1 for more details). Three successive steps for the emergence of *S*_*n*+1_ are represented, separated by black arrows. Blue and brown fonts are used to indicate the local and global number of *S*-haplotypes, respectively. Scenario **A** (local diversification): the compensatory mutation appears in the same deme as the initial mutation and the ancestral SC haplotype is still present in that deme.Scenario **B** (local replacement): the compensatory mutation appears in the same deme as the initial mutation when the ancestral SC haplotype is no longer present in that deme. Scenario **C** (allodiversification): the compensatory mutation appears in a different deme as the initial mutation and the ancestral SC haplotype is still present in that new deme. Scenario **D** (alloreplacement): the compensatory mutation appears in a different deme as the initial mutation and the ancestral SC haplotype is no longer present in that new deme. Scenario **E** (global replacement): the compensatory mutation appears in a different deme as the initial mutation and the ancestral SC haplotype is no longer present in the whole metapopulation. Note that the global numberof S-haplotypesincreasesonlyinscenariosA, B, CandD.

The first four scenarios lead to a net global diversification (an increase in the number of *S*-haplotypes at the global scale; Fig. 2A, 2B, 2C and 2D) but not the *Global replacement* scenario where the new *S*-haplotype replaces its ancestor in the whole metapopulation (Fig. 2E).

To evaluate the relative frequency of these five mutually exclusive scenarios in the dynamics of allelic diversification in a metapopulation we recorded the fate of every new SC haplotype during 1,000,000 generations in the 2-steps mutation model. We recorded the demes where the initial and the compensatory mutation occurred and the frequency of the ancestral *S*-haplotype in each deme when the compensatory mutation occurred.

## Results

In this study we asked three main questions: 1) does population structure affect the probability of diversification? 2) how does population structure affect the number of *S*-haplotypes relative to an isolated population, and if so, 3) what is the role of the 2-genes genetic architecture in determining these effects?

### Population structure decreases the overall probability of *S*-haplotype diversification

We first ran a hundred replicates of simulation runs for varying parameter sets under a 2-steps mutational model, starting with an initial number of *n* = 5 *S*-haplotypes. Decreasing the pollen dispersal rate *d*_*p*_ and hence increasing population subdivision decreased the parameter regions where diversification was possible, and decreased the overall probability of diversification (Fig. 3). For instance, for *µ* = 5 *×* 10^−6^, the parameter region of diversification was halved between *d*_*p*_ = 0.8 and *d*_*p*_ = 10^−5^. Increasing the mutation rate *µ*, expanded the parameter region of diversification (because more SC haplotypes were generated). Increasing the mutation rate to *µ* = 5 × 10^−3^, the effect of subdivision was much more subtle since the parameter region of diversification decreased only by *≈* 17% between *d*_*p*_ = 0.8 and *d*_*p*_ = 10^−5^. For the lowest value of the mutation rate that we explored (*µ* = 5 × 10^−7^), increasing the pollen dispersal rate first expanded the parameter region allowing diversification and then shrunk it. This non-monotonous effect of population structure could be an artifact of the cut-off number of generations after which we determined the state of the population, as the mutation-selection-drift equilibrium takes longer to be reached for low mutation rates. In most cases, the differences between the less structured population (*d*_*p*_ = 0.8) and the most structured population (*d*_*p*_ = 10^−8^) in the size of the parameter range allowing diversification were quantitatively similar to the differences between large and small non-structured populations (*N* = 5000 *vs. N* = 1000; Fig. 3). Thus, we hypothesized that the shrinkage of the diversification region with population structure is largely due to a reduction of the overall effective population size caused by the reduced pollen dispersal between demes.

**Fig. 3:**
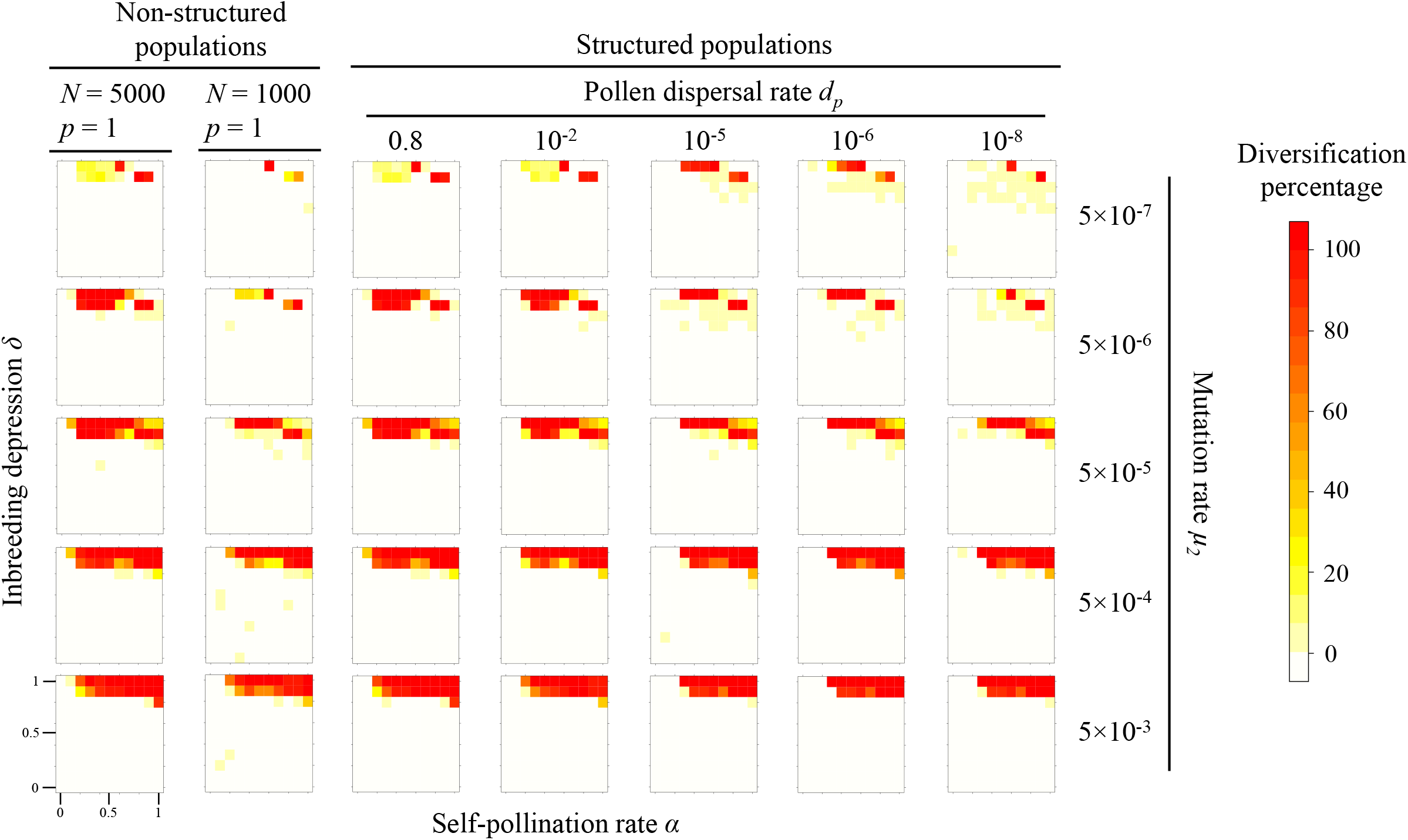
Diversification of *S*-haplotypes as a function of the mutation rate *µ* and the pollen dispersal rate *d*_*p*_. The metapopulation is composed of *p* = 5 demes containing 1,000 individuals each, except the two leftmost columns where the population is non-structured: a single deme of size 1,000 or 5,000. Color scaling on the right refers to the percentage of simulation runs among 100 replicates, for a given parameters set, where the total number of *S*-haplotypes increased in the metapopulation (starting with an initial number *n* = 5).

### Parallel local replacements in different demes promote global diversification

In order to study the effect of population structure on the number of *S*-haplotypes maintained in the metapopulation, we explored parameter values where the diversification frequency was higher than 50% for three values of the mutation rate *µ* and four values of the pollen dispersal rate *d*_*p*_. (Fig. S5). We found that the average number of *S*-haplotypes at the metapopulation level at steady state 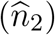 always increased with decreasing pollen migration rate *d*_*p*_ (Fig. 4 and Fig. S6). The magnitude of the increase ranged from 5% to 92% depending on the combination of parameters. The effect was especially marked for very low dispersal rates (*d*_*p*_ = 10^−8^) and was more subtle for higher dispersal rates (*d*_*p*_ = 10^−2^ and *d*_*p*_ = 10^−5^). Our results thus suggest that population subdivision favors the maintenance of a higher *S*-haplotype diversity, but mostly in strongly structured metapopulations.

**Fig. 4:**
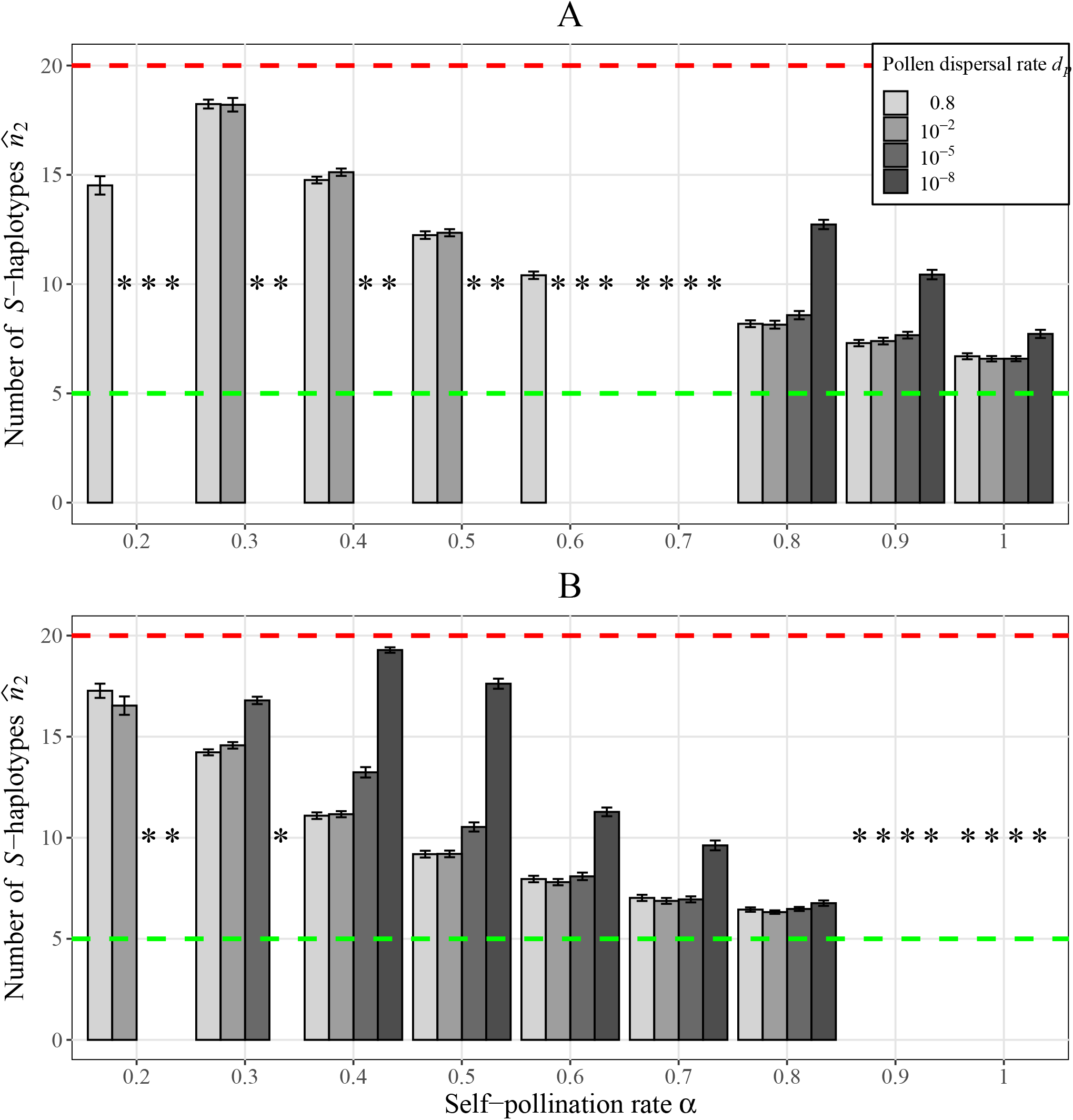
Average number and 95% CI of *S*-haplotypes at steady state as a function of the self-pollination rate *α* and for different pollen dispersal rates *d*_*p*_. An asterisk indicates a combination of parameters where diversification is too rare to be considered (inferior to 50% of diversification *i*.*e*. regions surrounded in black in Fig. S5). Dashed green and red line show the initial number of *S*-haplotypes *n* and the maximum number of *S*-haplotypes *k*, respectively. Parameters:*µ*_2_ = 5 × 10^−5^, *N* = 5000, *p* = 5, *n* = 5 and *k* = 20. A: *δ* = 0.9; B: *δ* = 1.

We then aimed at describing how the two successive mutations that are necessary for the formation of new haplotypes occurred. We recorded whether the second (compensatory) mutation appears in the same or in a different deme than the ancestral *S*-haplotype, and whether the ancestral *S*-haplotype was lost at the local (deme) or the global (metapopulation) scale (Fig. 5). The relative importance of the different fitness valley crossing scenarios differed sharply between largely unstructured (*d*_*p*_ = 0.8) and structured populations (*d*_*p*_ = 10^−5^ and 10^−8^) (Fig. 5). In unstructured populations (*d*_*p*_ = 0.8), when the self-pollination rate was low (*α ≤* 0.6), the emergence of new *S*-haplotypes was most of the time accompanied with global loss of the ancestral *S*-haplotype, with successive mutations occurring in different demes (global replacement). With higher self-pollination rates in unstructured populations, global replacement became as frequent as allodiversification (*i*.*e*. the scenario in which the two mutations arise in different demes but the ancestral *S*-haplotype is conserved). This is due to the fact that SC haplotypes can more easily exclude their ancestral haplotype when they are still able to frequently outcross. The emergence of new *S*-haplotypes by succession of the two mutations in the same deme (either local replacement or local diversification) accounted for a little less that 25% of the formations of new *S*-haplotypes for all parameter values explored, *i*.*e*. in 75% of the cases, the first and second mutations occurred in different demes. In highly structured populations, in contrast, new *S*-haplotypes arose almost exclusively through within-deme events (local replacement and local diversification), and global replacements were rare. Our results thus show that population structure increases global *S*-haplotype diversity both because diversification and replacement take place in parallel in different demes, and because the loss of ancestral *S*-haplotypes at the scale of the metapopulation also becomes more rare.

**Fig. 5:**
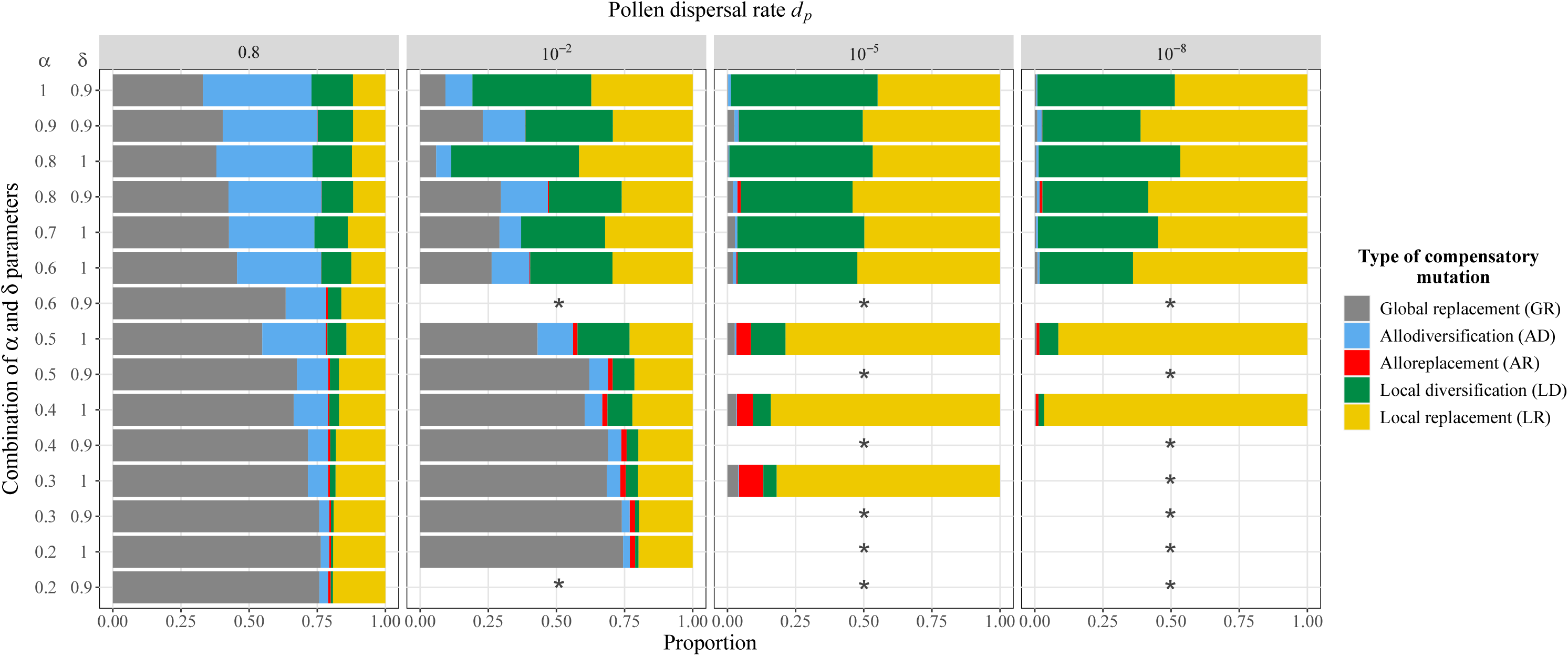
Proportion of five possible scenarios for the emergence of new *S*-haplotypes in a metapopulation over 100 simulation runs (see Fig. 2). An asterisk indicates a combination of parameters where diversification is too rare to be considered (inferior to 50% of diversification *i*.*e*. regions surrounded in black in Fig. S5). Parameters: *µ*_2_ = 5 × 10^−5^, *N* = 5000, *p* = 5, *n* = 5 and *k* = 20.

### Interaction between population subdivision and genetic architecture increases *S*-haplotype diversity

Our last goal was to check whether the genetic architecture underlying SI could change the effect of population structure on *S*-haplotype diversity. To achieve this, we compared the average *S*-haplotype diversity in structured populations under the 1- *vs*. 2-steps mutational models. We determined the (effective) mutation rate of the 1-step mutational model under which the average number of *S*-haplotypes best approximated that under the 2-steps mutational model 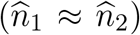 in an unstructured population. For all parameter sets we studied, we found that the 2-steps mutational model was more sensitive to population structure, consistently producing new *S*-haplotypes at a higher rate than the 1-step mutational model (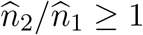 in Fig. 6). The contrast between the two mutational models was higher when the dispersal rate decreased, resulting in a higher number of *S*-haplotypes under the 2-steps model than under the 1-step model. The number of *S*-haplotypes can be twice higher under the 2-steps than under the 1-step mutational model, even though our simulations were calibrated so that the two mutation models behaved identically in unstructured populations. Our results thus show that in structured populations, a more complex genetic architecture underlying SI can favor the emergence of *S*-haplotype diversity even when controlling for differences in the effective mutation rates.

**Fig. 6:**
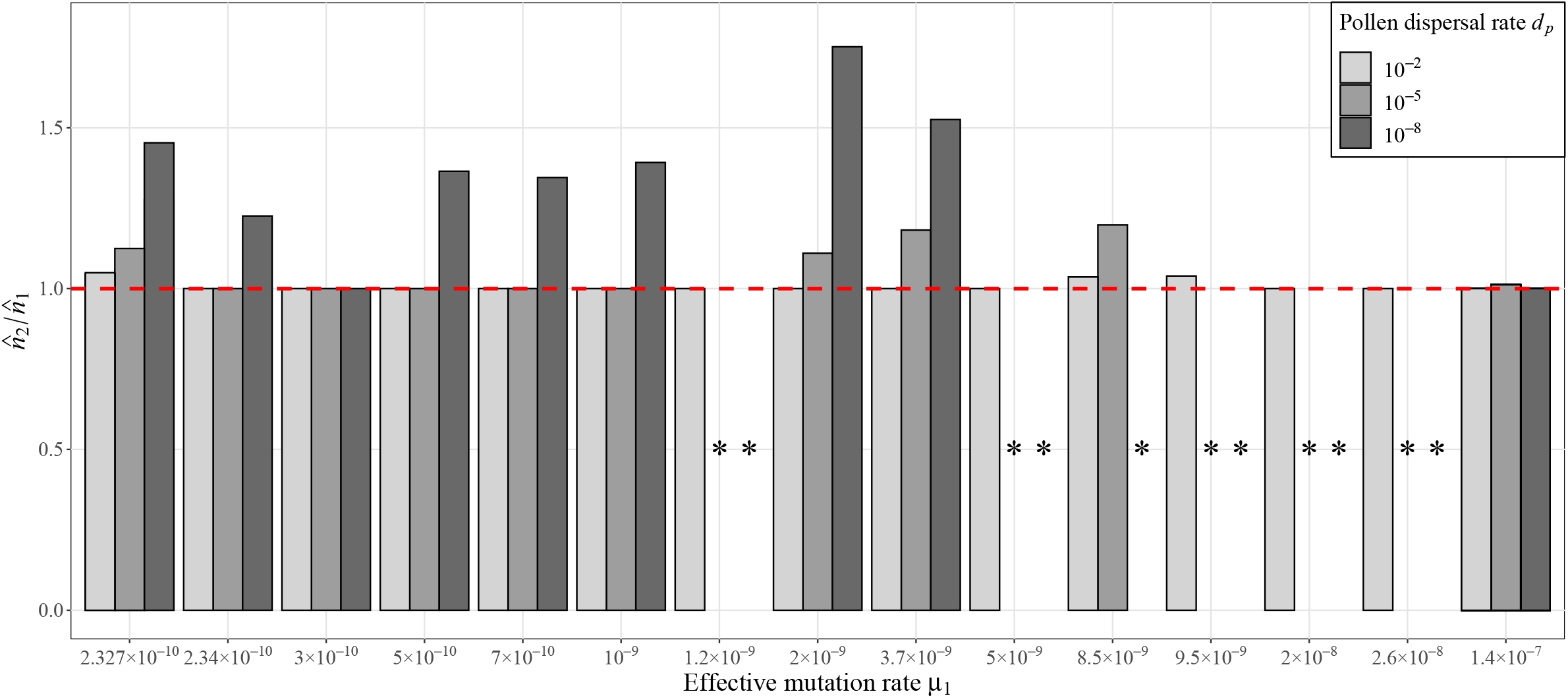
Ratio of the number of *S*-haplotypes at steady state between the 2-steps *vs*. 1-step mutation models 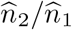 for a given effective mutation rate *µ*_1_ (see text for details). The red dashed line represents a ratio of 1 (with a 95% CI): in other words bars higher than the red dashed line mean that the number of *S*-haplotypes at equilibrium is significantly higher under the 2-steps than the 1-step mutation model even though the effective mutation rate is similar. An asterisk indicates a combination of parameters where diversification is too rare to be considered (inferior to 50% of diversification *i*.*e*. regions surrounded in black in Fig. S5).

## Discussion

### Population structure decreases the conditions under which diversification occurs, yet increases the number of *S*-haplotypes maintained

Our results show that population structure monotonically increases the *S*-haplotype diversity at the metapopulation level when diversification was possible, although it also decreases the probability of *S*-haplotype diversification overall. It has been shown, in a 1-step mutation model, that the *S*-allele diversity at the metapopulation level varies non-monotonically with subdivision (Schierup, 1998; Muirhead, 2001). Indeed, subdivision increases the loss of local *S*-alleles by drift and this loss is compensated by the differentiation of haplotypes among demes only at low migration rates (of the order of the mutation rate) because balancing selection tends to homogenize *S*-allele diversity among demes. In the 2-steps mutation model, our results show that limited dispersal enhances global diversity. While the same phenomena at the local scale also apply, this genetic architecture modifies the interactions between the underlying mechanisms. Indeed, even after controlling for different effective mutations rates under the 1- *vs*. 2-steps mutation models, *S*-haplotype diversity was always higher under the 2-steps than the 1-step mutation model (Fig. 6). This suggests that crossing fitness valleys through partial self-fertilization, as implied by the 2-steps model, modifies the interactions between selection and dispersal, ultimately affecting the diversity of *S*-haplotypes.

Two mechanisms can explain why population structure increases the *S*-haplotype diversity. First, as shown by Gervais et al. (2011), the diversification rate is higher when the number of haplotypes is low. Since the relative fitness of SC haplotypes (compared to *S*-haplotypes) is higher when the number of *S*-haplotypes is low, their expected frequency is also higher, which makes the appearance of a compensatory mutation more probable. Population structure thus enhances *S*-haplotype diversity because locally the number of *S*-haplotypes is decreased, causing diversification to be faster at the local scale. This is supported by our observations that the occurrence of local diversification events increased with decreasing dispersal. Second, the relative importance of scenarios by which fitness valleys were crossed varies with population structure. In a non-structured population, new *S*-haplotypes emerge either by a replacement or a diversification (Gervais et al., 2011). In a structured population, global diversification also comes from events of local replacement. As replacements taking place in different demes likely affects different *S*-haplotypes, different *S*-haplotypes will be lost and replaced locally. Replacement occurring in different demes thus increases *S*-haplotype diversity at the global scale. These two mechanisms probably compensate for the homogenization of *S*-haplotype diversity among demes caused by balancing selection, explaining why the number of *S*-haplotypes increases monotonically with subdivision rather than non-monotonically as in the 1-step mutation model (Schierup, 1998; Muirhead, 2001).

Overall, can limited dispersal help fill the gap between the observed and expected *S*-haplotype diversity? While we do see an increase in the number of *S*-haplotypes caused by population structure, even for very small dispersal rates, the increase is quantitatively moderate, and still far from explaining the very large *S*-haplotype diversity observed in natural populations (from a few dozens to *>*100 haplotypes, Lawrence 2000). An obvious potential caveat is that the number of *S*-haplotypes that can be obtained in our model is limited by the assumed mutation process. Indeed, for compensatory mutations to emerge, one has to set a maximum number of alleles at each locus (our *k* parameter), imposing a maximum number of *S*-haplotypes. Gervais et al. (2011) showed that decreasing *k* increases the steady-state number of *S*-haplotypes 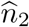 because it increases the probability of compensatory mutation restoring SI in an SC haplotype (Fig. S3). Yet, in our simulations, the steady state number of *S*-haplotypes is generally much lower than the theoretical maximum (Fig. 4), suggesting that crossing fitness valleys is more affected by population structure and selective mechanisms than the mutational process.

Since population isolation tends to enhance the local diversification dynamics, the large diversity in natural populations might be the result of the complex demographic history of SI species, where populations have several times fragmented and merged, expanded and shrunk, lost and gained pollinators, or exchanged *S*-haplotypes with closely related species. It may also be that other mechanisms, not taken into account in the present model, contribute to the rise of new *S*-haplotypes in nature. For example, Sakai (2016) proposed that *S*-haplotypes can have been generated from SC populations by progressive optimization of the self-recognition capacity between *S*-haplotypes that initially have incomplete recognition and rejection. He found that this model can generate up to 40-50 haplotypes in an unstructured SC population within a few hundred generations, suggesting that the large *S*-haplotype diversity is the relic of the emergence of SI. How population structure would affect *S*-haplotype diversity in this model is an open question.

### Fitness valley crossing in structured populations under balancing selection

The crossing of fitness valleys has been studied by population genetics models under directional selection, where the fitness landscape is static as it results from fixed parameters. Balancing selection makes the fitness landscape variable because its shape depends on the frequency of the different haplotypes. Fitness valley crossings under directional selection can occur through two mechanisms (Weissman et al., 2009): by stochastic tunnelling, where intermediate deleterious genotypes segregate at low frequency because of recurrent mutations, or by sequential fixation, where the intermediate genotype first gets fixed and then the advantageous genotype arises by mutation and goes to fixation. Population structure can facilitate the crossing of fitness valleys under directional selection, a phenomenon interpreted as resulting from “competitive assortment” (Bitbol and Schwab, 2014; Komarova et al., 2014; Cooper and Kerr, 2016; McLaren, 2016). As dispersal is limited and competition between genotypes occurs locally, local kinship decreases the relative disadvantage of intermediate deleterious genotypes relatively to their wild-type counterparts. The frequency of intermediate genotypes is thus higher, they can even possibly get fixed locally, which facilitates stochastic tunnelling and sequential fixation.

In our analysis, the temporary segregation of SC intermediate haplotypes in local demes due to limited dispersal can be interpreted as competitive assortment and may thus facilitate *S*-haplotype diversification through stochastic tunnelling. Diversification is expected to be proportional to the residence time of SC intermediate haplotypes within demes, because the longer the residence time, the higher the probability that a compensatory mutation will occur. The residence time of SC intermediate haplotypes depends on selection acting on SC intermediate haplotypes (depending on the relative importance of inbreeding depression and mate availability). Competitive assortment mitigates the effect of selection against SC intermediate haplotypes because the mate availability advantage increases when the local kinship is higher (a higher kinship means a lower number of *S*-haplotypes). In contrast, local sequential fixation (sequential fixation in a single deme) is unlikely to be common. Indeed, the coexistence of a deme where a SC variant is fixed along with other demes where SI is conserved is never observed in a finite islands population (see also Brom et al., 2020). This is because, neglecting stochastic fixation, if the conditions are met for a SC haplotype to fix in one deme, SC haplotypes will also tend to get fixed in all other demes of the metapopulation. Overall, fitness valley crossing in the case of *S*-haplotypes diversification in a subdivided population is mostly due to stochastic tunnelling rather than sequential fixation.

A series of models (Bod’ová et al., 2018; Harkness et al., 2019; Harkness and Brandvain, 2021) have focused on a deeply rooted SI system in Angiosperms based on non-self recognition of the pollen (Fujii et al., 2016). The mutational pathway underlying the emergence of new *S*-haplotypes in such systems is different from the one we supposed here. Bod’ová et al. (2018) found that in such a system new *S*-haplotypes can emerge through a variety of pathways, in particular frequently by a SC intermediate haplotype (*i*.*e*. a fitness valley crossing). The SC intermediate pathway becomes more likely compared to other pathways when the number of specificities increases and when the mutation rate decreases. How population structure affects diversity in non-self recognition SI systems is another interesting open question.

### Relevance to the study of natural populations

Overall, our model suggests that the residence time of SC intermediate haplotypes is essential for *S*-haplotype diversification. We can speculate that new *S*-haplotypes are most likely to occur in small isolated demes, or in populations at the margins of species range where SI is most frequently lost *e*.*g. Arabidopsis lyrata* (Griffin and Willi, 2014). Observations in such populations are now needed to test the models’ predictions.

It is not yet totally understood why some species have lost SI while others have not, even among sister species. It is possible that different demographic histories, including histories of habitat fragmentation and pollinators loss may lead to these different outcomes. Indeed, Brom et al. (2020) predicted that decreasing dispersal rate in a subdivided population should facilitate the loss of SI because the local effective number of *S*-haplotypes is decreased. Encinas-Viso et al. (2020) further showed that the loss of SI can also occur during range expansion in the range margins and extend to the core population under certain conditions. However, here, we showed that population subdivision makes *S*-haplotype diversification less likely yet faster when it occurs. Hence, we argue that the evolutionary dynamics of a SI species during range expansion, or which recently suffered a fragmentation or pollinators loss is bistable. Either SI will be lost in the short run, or SI will diversify again rapidly. In a nutshell, the loss of SI and SI diversification can be viewed as conflicting processes, and the evolutionary trajectory followed by a species is the result of the race between both processes. A better understanding of how demographic processes and genetic mechanisms affect the balance between SI diversification and SI loss could help predict whether SI loss or SI rescue is the most likely scenario after range expansion, fragmentation, population bottleneck, or the loss of a pollinator.

## Acknowledgements

This work was funded by the European Research Council (NOVEL project, grant #648321). The authors also thank the Région Hauts-de-France, the Ministère de l’Enseignement Supérieur et de la Recherche (CPER Climibio), and the European Fund for Regional Economic Development for their financial support.

---

**Fig. S1.**
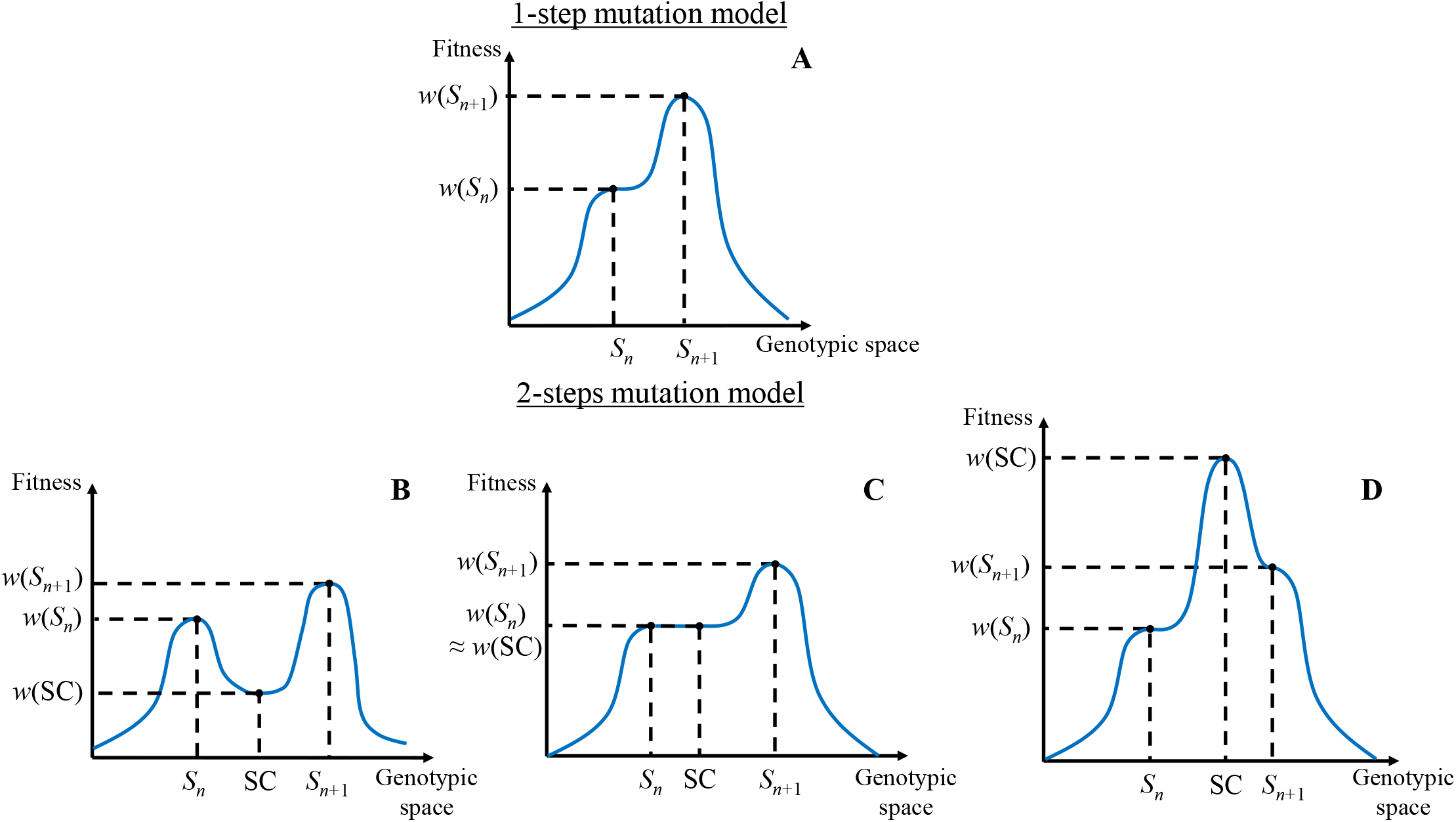
Fitness landscapes when a new *S*-haplotype *S*_*n*+1_ emerges under a 1-step or (**B, C, D**) a 2-steps mutation model. X-axis: simplified 1D genotypic space where: *S*_*n*_ represents the *n S*-haplotypes in the population before the emergence of the new *S*-haplotype *S*_*n*+1_; and *S*_*b*_ represents a self-compatible intermediate haplotype in a 2-steps mutation model (see text for further details). Y-axis: relative fitness of each haplotype assuming that a new *S*-haplotype *S*_*n*+1_ emerges at low frequency into a resident population, at mutation-drift-selection equilibrium, composed of *n S*-haplotypes with identical frequencies and a self-compatible intermediate haplotype. Because of negative frequency dependent selection, *S*_*n*+1_ has a higher fitness on the landscape than the other *S*_*n*_ haplotypes. The self-compatible intermediate haplotype can have three different positions on the fitness landscape: (**B**) it has either a fitness lower than all *S*-haplotypes; or it is present in the population because of recurrent mutation and/or genetic drift; or the emergence of a new *S*-haplotype *S*_*n*+1_ needs the crossing of a fitness valley (stochastic tunneling); (**C**) The self-compatible intermediate haplotype coexists with *S*_*n*_ haplotypes at equilibrium (they have the same fitness); the emergence of the new *S*-haplotype *S*_*n*+1_ needs the crossing of a flat landscape; (**D**) The relative fitness of the self-compatible intermediate haplo2ty9pe is higher than all *S*-haplotypes (including the new emerging *S*_*n*+1_); the self-compatible intermediate haplotype will eventually go to fixation and the self-incompatibility system will be lost.

**Fig. S2.**
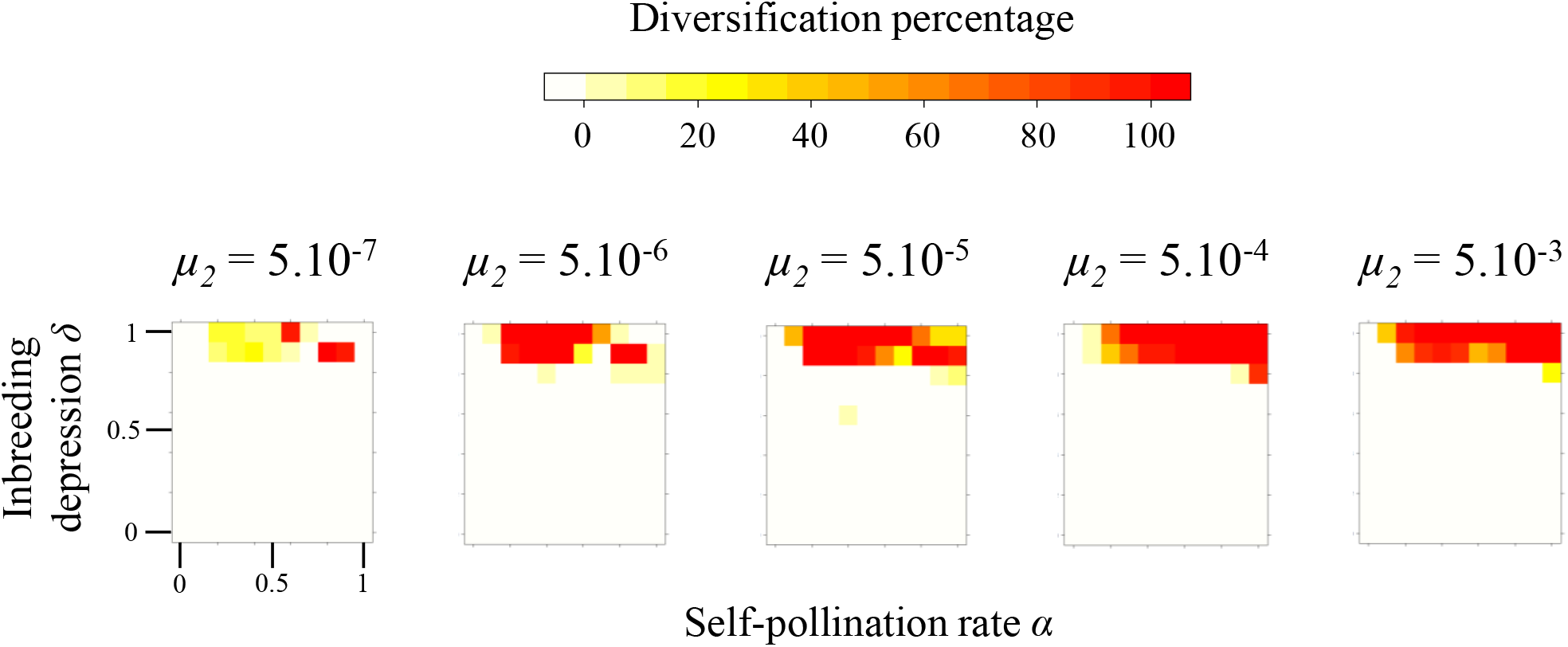
Parameter regions allowing diversification of *S*-haplotypes — run with our simulation program of the 2-steps mutation model — as a function of the mutation rate *µ*_2_ for a non-structured population. The colour range indicates the diversification percentage (the number of times diversification occurred over 100 replicates) and goes from white, meaning no diversification occurred, to dark red, meaning diversification always occurred. Each grid cell corresponds to 100 replicates. For all simulations *n* = 5, *k* = 20, *N* = 5000, *p* = 5.

**Fig. S3.**
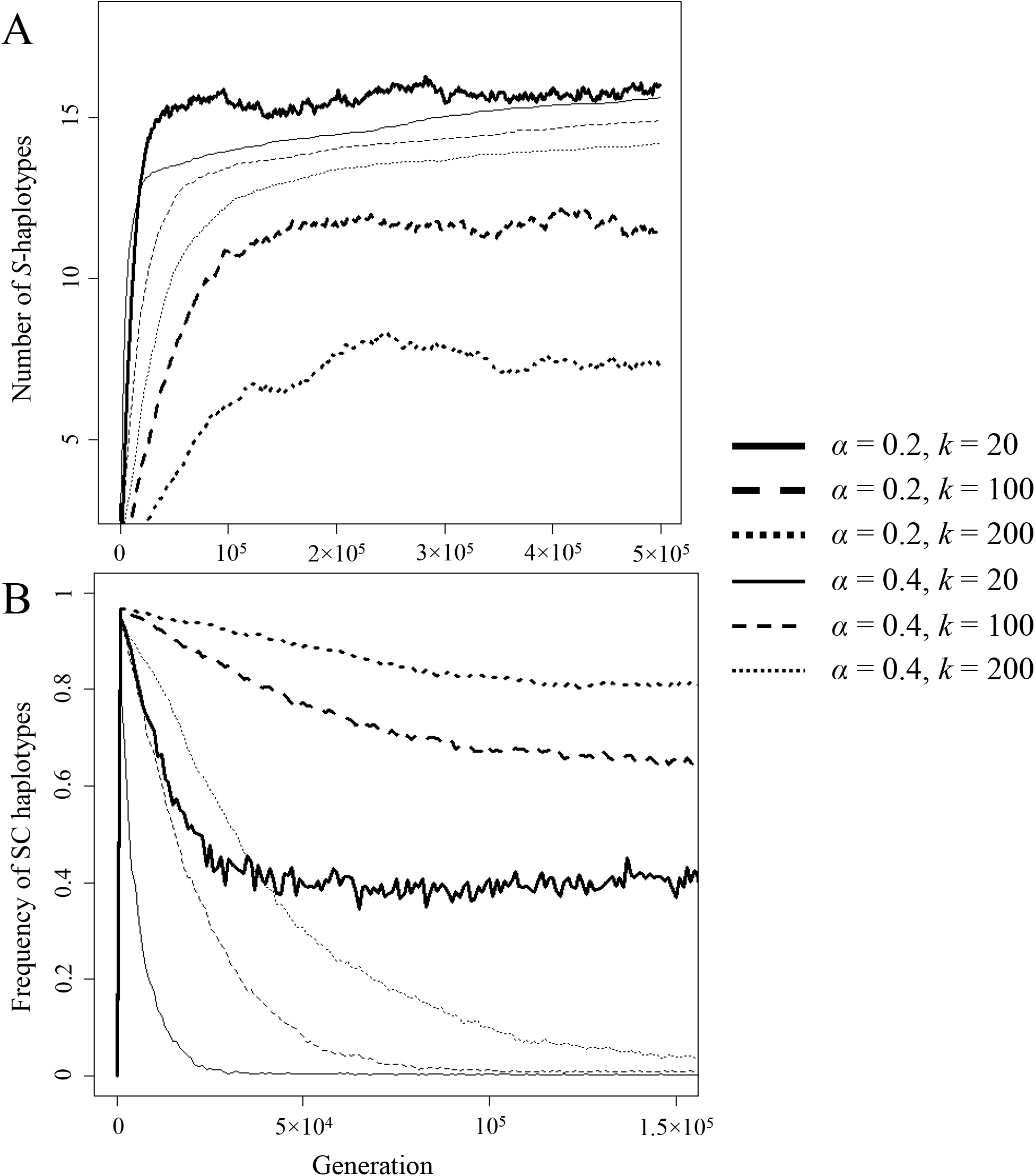
Dynamics of the number of haplotypes for different combinations of self-pollination rate *α* and number of specificities *k*. Thick line: *α* = 0.2, thin line: *α* = 0.4, continuous line: *k* = 20, dashed line: *k* = 100, dotted line: *k* = 200. Dynamics are averages over 100 replicates and were calculated for *δ* = 0.9, *N* = 5, 000, *p* = 5 and *n* = 5. **A**. Number of *S*-haplotypes across 5 × 10^5^ generations. **B**. Frequency of SC haplotypes across 1.5 × 10^5^ generations.

**Fig. S4.**
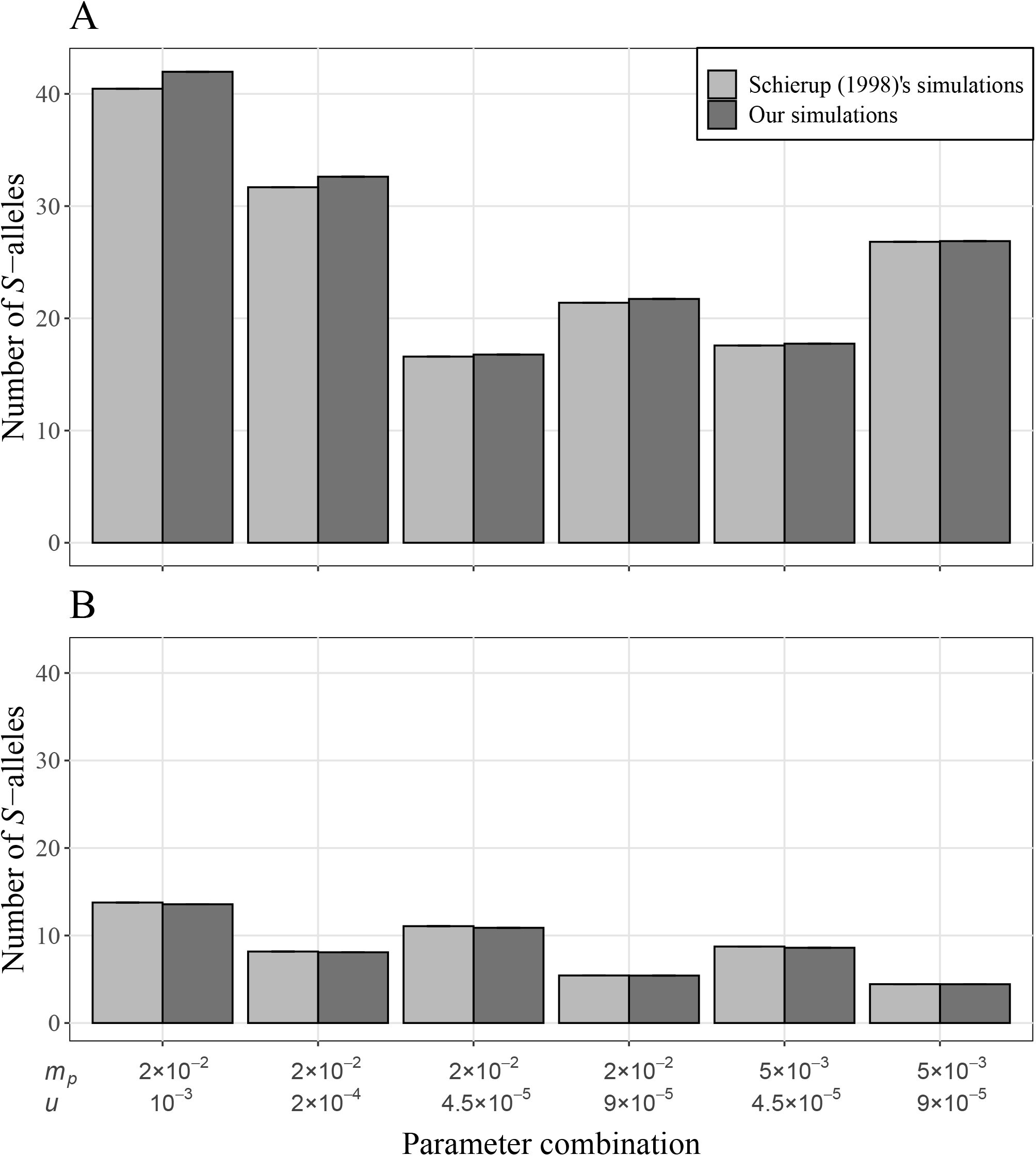
Verification that the number of *S*-alleles in the population (**A**) and average number of alleles per deme (**B**) calculated with our simulation program (light-grey bars) matches those in Schierup (1998) (dark-grey bars) with the method detailed in Schierup (1998) for different combinations of parameters (pollen migration rate *m*_*p*_ or *d*_*p*_ and mutation rate *u* or *µ*_1_). Each bar represents the average number of alleles over 100 replicates and the interval *±*SE is represented by error bars (often as thick as the upper line of the bar).

**Fig. S5.**
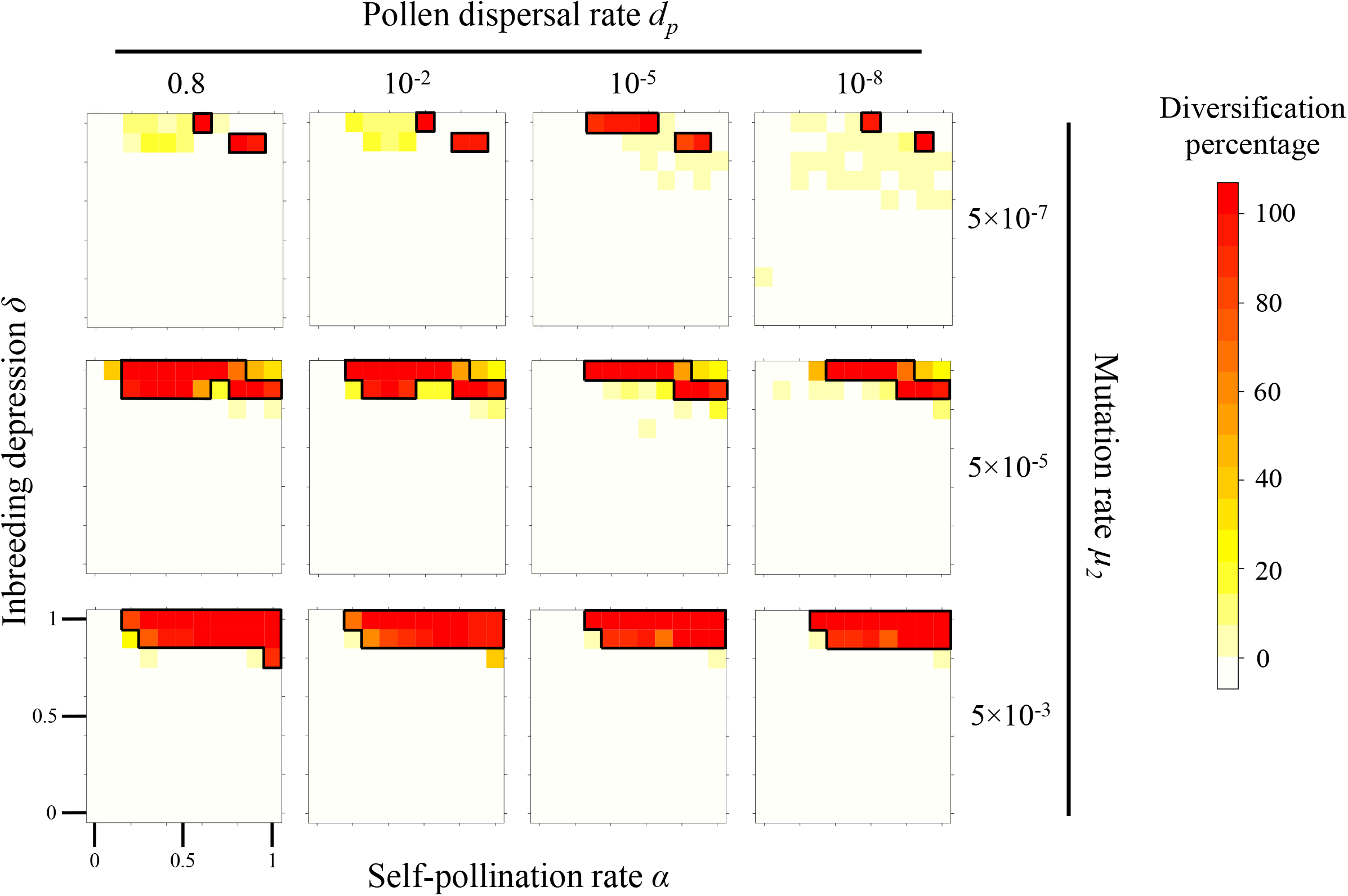
Parameter regions allowing diversification of *S*-haplotypes as a function of mutation rate *µ* and pollen dispersal rate *d*_*p*_. The colour range indicates the diversification percentage (the number of times diversification occurred over 100 replicates) and goes from white, meaning no diversification occurred, to dark red, meaning diversification always occurred. Regions surrounded in black correspond to combination of parameters simulated to study the haplotype number at equilibrium 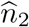 (Fig. 4). Each grid cell corresponds to 100 replicates. For all simulations *N* = 5000, *p* = 5, *n* = 5 and *k* = 20.

**Fig. S6.**
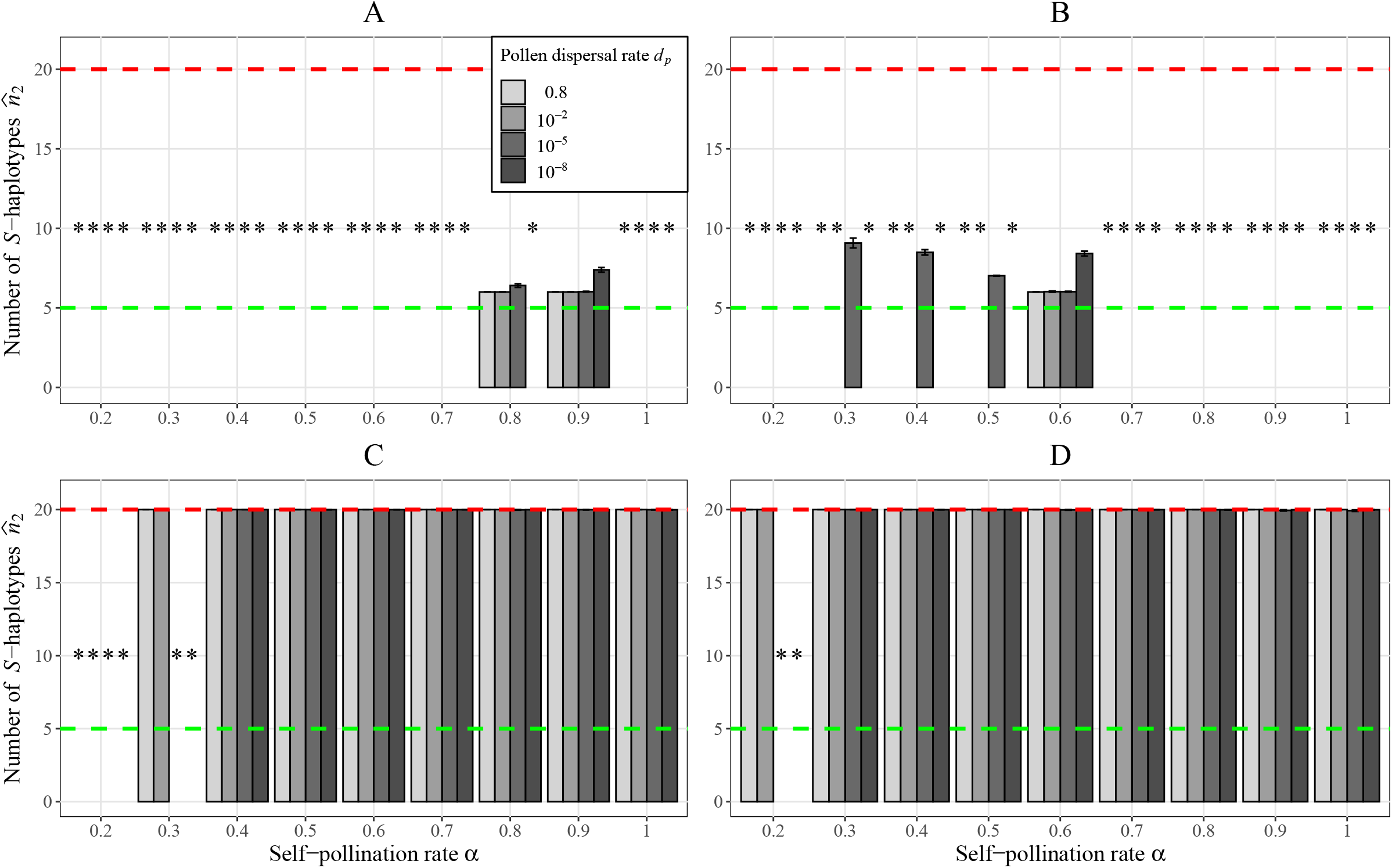
Average number of *S*-haplotypes at steady state 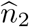 and 95% CI (not visible when too small) as a function of the self-pollination rate *α* and for different pollen dispersal rate *d*_*p*_. An asterisk indicates a combination of parameters where diversification is too rare to be considered (see text for details, and see Fig. 3). Dashed green and red line show the initial number of *S*-haplotypes *n* and the maximum number of *S*-haplotypes *k*, respectively. Parameters: *N* = 5000, *p* = 5, *n* = 5 and *k* = 20. A: *δ* = 0.9, *µ*_2_ = 5 × 10^−7^; B: *δ* = 1, *µ*_2_ = 5 × 10^−7^; C: *δ* = 0.9, *µ*_2_ = 5 × 10^−3^; D: *δ* = 1, *µ*_2_ = 5 × 10^−3^.

